# Uncovering the Molecular Regulation of Seed Development and Germination in Endangered Legume *Paubrasilia echinata* Through Proteomic and Polyamine Analyses

**DOI:** 10.1101/2025.10.13.682162

**Authors:** Rosana Gobbi Vettorazzi, Renan Carrari-Santos, Kariane Rodrigues Sousa, Tadeu dos Reis de Oliveira, Clicia Grativol, Geovanna Vitória Olimpio, Thiago Motta Venancio, Vitor Batista Pinto, Gabriel Quintanilha-Peixoto, Vanildo Silveira, Claudete Santa-Catarina

## Abstract

Understanding the molecular regulation of seed maturation and germination is essential for plant conservation and agricultural applications. Here, we provide novel insights into the proteomic and polyamine dynamics of seeds from *Paubrasilia echinata*, an endangered legume, at 4, 6, and 8 weeks after anthesis. Using a sequential protein extraction approach combined with a species-specific protein database, we identified over 2,000 proteins, uncovering key regulators of maturation, stress tolerance, and germination. Seeds that reached maturity at 6 weeks displayed a high accumulation of proteasome components, translation machinery, and stress-associated proteins, notably late embryogenesis abundant (LEA) and heat shock proteins (HSPs), underscoring their role in seed development. Polyamine profiling showed that high putrescine levels were associated with early development and reduced germination, while spermidine and spermine were correlated with maturation, tolerance to desiccation, and germination. The decreases in certain proteins and polyamines during the transition to germination reveal dynamic regulatory shifts essential for the establishment of seedlings. These molecular signatures reveal dynamic metabolic and regulatory changes that facilitate the transition from seed development to germination. These findings reveal previously uncharacterized mechanisms of seed maturation and early germination in *P. echinata* and provide information on seed biology, ex situ conservation, and propagation strategies for endangered legumes.

## 1. Introduction

Known as brazilwood (Pau-brasil), *Paubrasilia echinata* (Lam.) Gagnon, H.C. Lima & G.P. Lewis (Fabaceae) is a native tree species of the Brazilian Atlantic Forest that is of great biological, historical, and cultural importance to Brazil and is officially recognized as a national tree.^1,2^ This species has been exploited since its colonization for dye extraction and later for the manufacture of high-quality violin bows.^3–6^ Owing to its history of exploitation, *P. echinata* is classified as endangered and has been listed on the CITES^7^, the IUCN Red List,^8^ the Red Book of the Brazilian Flora,^4^ and the Official List of Brazilian Flora Species Threatened with Extinction.^9^

The management of *P. echinata* seeds presents challenges, such as irregular fruiting, a high incidence of seed/ovule abortion, seed predation, and rapid loss of germination potential, which hinder the recovery of native populations.^10–12^ Furthermore, there is limited information is available regarding the biochemical aspects of seed development and germination in this species.^10,13^ Among these studies, Borges et al.^14^ evaluated variations in sugar and cyclitol contents, whereas Leduc et al.^15^ and Mescia et al.^16^ investigated the carbohydrates involved in the induction of tolerance to desiccation during seed maturation. This lack of basic knowledge about seed behavior hampers the implementation of conservation strategies and propagation techniques for forest species.

Research in biochemical and molecular biology enables the identification of regulators necessary in zygotic embryogenesis, seed development, and germination processes.^17–19^ These studies can also be applied to optimize somatic embryogenesis, as somatic and zygotic embryos exhibit analogous development patterns.^20^ In recent decades, important advances have been made in high-throughput technologies such as RNA sequencing (RNA-seq) and proteomics. Integrating RNA-seq data with proteomics data is a powerful strategy to expand and refine protein identification.^21^ To date, there are no publicly available transcriptomic data for *P. echinata*. This raises the question of why such a valuable species is so poorly studied and makes it essential to generate more genetic sequence information to study the species further. Sequencing technology provides a rapid and efficient approach to generate transcriptome data from non-model organisms that do not have a complete genome sequence.^22,23^ Compared with whole genome sequencing, RNA sequencing (RNA-seq) is low-cost and high-throughput; therefore, it is an important approach in research and may constitute a strategy to improve proteomic analyses in non-model organisms.^22–24^

In recent decades, proteomics has emerged as a powerful approach for investigating protein functions and interactions in complex biological systems.^25^ Proteins are responsible for various metabolic processes in seeds and serve as important structural components in the cytoskeleton, membranes, cell wall, and more.^26,27^ Proteomic studies enable the identification of differentially accumulated proteins (DAPs) under different conditions and have been applied in seed science and technology,^28^ particularly for studying seed development and germination in woody species, including *Cariniana legalis*,^29,30^ *Quercus ilex*,^31^ and *Araucaria angustifolia*.^32^ These studies offer valuable insights into the protein dynamics essential for germination.

In addition to the proteomic profile, changes in the content of PAs such as putrescine (Put), spermidine (Spd), and spermine (Spm) during seed development and germination have also been studied in several woody species, including *Ocotea catharinensis*,^17^ *Cedrela fissilis*,^18^ *C. legalis*,^30^ *Plathymenia foliolosa,* and *Dalbergia nigra*.^33^ PAs are low-molecular-weight aliphatic compounds that can interact with macromolecules, including DNA, RNA, proteins, phospholipids, and components of the cell wall, through hydrogen and ionic bonding.^34,35^ PAs also regulate cell division, differentiation, and gene expression, as well as the synthesis of several proteins at the translational level, and play important roles in seed development, germination, and seedling growth.^36–39^

The patterns of protein and PA accumulation vary among species. Despite advances in research, studies on the development and germination of woody tree seeds are still very limited, and the relationships between proteomic profiles and PA content during the development and germination of *P. echinata* seeds remain unexplored. This study aimed to investigate the associations between proteomic profiles and PA content throughout the maturation and germination stages of *P. echinata* seeds. This research contributes to the development of improved conservation and propagation strategies for this species.

## 2. Materials and Methods

### 2.1. Plant material

The development of the seeds was assessed using fruits containing immature seeds of *P. echinata* harvested 4 and 6 weeks after anthesis (WAA) and mature dispersed seeds at 8 WAA, which were collected in November and December 2019 from trees located in Campos dos Goytacazes – RJ (21 ° 45 ’59.1’S 41 ° 16’17.8’W). The samples were deposited at the Herbarium of the Universidade Estadual do Norte Fluminense Darcy Ribeiro (HUENF00009622). The seeds were extracted from the fruits and used for seed development and germination studies.

### 2.2. Determination of length, fresh matter (FM), and dry matter (DM)

The length (cm), FM, and DM of the seeds were measured using five biological replicates, each consisting of five seeds. The length was determined with a millimeter ruler. FM was obtained by weighing the seeds on an analytical balance (Shimadzu; Kyoto, Japan). The samples were then placed in paper bags and dried in an oven with forced air circulation at 70 ° C for 72 h to determine the DM.

### 2.3. Germination and Germination Speed Index (GSI)

The percentage of germination and GSI of the seeds at each collection time were assessed. Germination was performed according to Mello and Barbedo^40^ using four biological replicates, each with 25 seeds. Before germination, seeds collected at 4, 6, and 8 WAA were subjected to a single wash with water containing three drops of commercial detergent, followed by rinsing with distilled water. The seeds were subsequently placed on Germitest^®^ paper (J Prolab; Paraná, Brazil), moistened with type 2, deionized and autoclaved water in a ratio of 2.5 times the DM of the substrate. The paper rolls were enclosed in polyethylene bags and placed in BOD-type germination chambers (Eletrolab; São Paulo, Brazil) maintained at 25 ° C, at 8 h photoperiod, with a continuous light intensity of 40 µmol m^-2^ s^-1^.

For GSI analysis, daily observations were made for eight days to determine the number of germinated seeds while considering the protrusion of the radicle. GSI was calculated according to Maguire^41^, considering the normal seedlings obtained, that is, those with well-developed aerial parts and root systems capable of continued growth.^42^ After eight days, the FM and DM of the seedlings were measured.

### 2.4. Development of a non-redundant protein database of *P. echinata*

Protein identification was carried out using an in-house nonredundant protein database of *P. echinata* generated from high-throughput RNA sequencing (RNA-seq) data. For the generation of RNA- seq data, RNA extraction was performed on samples composed of immature seeds harvested at 6 WAA (I); dispersed seeds harvested at 8 WAA (II); dispersed seeds collected after the emergence of the radicle at 24 h of germination (III); 8-day seedlings grown at 25 ° C in the Petri dish with type 2 autoclavated, desionized water (IV); 8-day seedlings grown at 25 ° C under 4 mM FeSOC solution (V); 8-day seedlings grown at 25 ° C in the Petri dish with type 2 autoclavated, desionized water, then exposed to 40 °C for six h (VI); 8-day seedlings grown at 25 ° C in the Petri dish with type 2 autoclavated, deionized water, then kept for 6 h without water (VII); 8-day seedlings at 25 ° C in the Petri dish with type 2 autoclavated, deionized water, then kept for 2 h in the sun with the average photosynthetically active radiation (PAR) of 2.372,25 µmol mC² sC¹ (Fieldscout Quantum Light 6 Sensor Reader - Spectrum technologies, Aurora, USA) (VIII); 8-day-old seedlings at 25 °C in Petri dish with 6% PEG 3350 solution (IX); somatic embryos from globular to cotyledonary stages (X); leaves of 30-day-old seedlings *in vitro* germinated (XI); and roots of 30-day-old seedlings *in vitro* germinated (XII).

For RNA extraction, approximately 500 mg FM from each sample was flash frozen in liquid nitrogen and stored at −80 ° C. Initially, the samples were pulverized in liquid nitrogen using a ceramic mortar and pestle. The resulting powder from each sample was transferred to new tubes containing 1 mL of extraction solution consisting of 100% phenol, 100% chloroform, and extraction buffer (200 mM Tris- Cl treated with diethylpyrocarbonate (DEPC) (pH 7.5), 100 mM LiCl (treated with DEPC), 5 mM EDTA and 1% SDS) in a 1:1:2 ratio (v/v/v). The samples were then vortexed for 1 min followed by centrifugation for 10 min at 20,000 × g. The resulting aqueous phase was transferred to a new tube containing 500 μL of phenol and chloroform (1:1; v/v), vortexed for 1 min and centrifuged for 10 min at 20,000 × g (from this step, the material was always kept on ice). The resulting aqueous phase (∼500 μL) was transferred to a new tube and 500 μL of 6 M LiCl (DEPC-treated) was added. The tubes were incubated at 4 °C for 16 h, followed by centrifugation for 10 min at 20,000 × g. The supernatant was discarded and the precipitate was resuspended in 1 mL of 3 M LiCl (DEPC-treated) followed by centrifugation for 10 min at 20,000 × g at 4 °C. The supernatant was discarded and the precipitate was resuspended in 250 μL of DEPC-treated H_2_O. Then, 25 μL of 3 M NaOAc (treated with DEPC) and 550 μL of absolute ice cold ethanol were added. The tubes were incubated for 30 min at –70 °C and centrifuged for 15 min at 20,000 × g at 4 °C, after which the supernatant was discarded. The precipitate was carefully washed with ∼ 300 μL of 70% ethanol (DEPC treated), followed by centrifugation for 15 min at 20,000 × g at 4 °C. Subsequently, the ethanol was discarded and the RNA was resuspended in 50 μL of DEPC-treated H_2_O. The RNA concentration of each sample was determined with a NanoDrop 2000c spectrophotometer (Thermo Scientific; Wilmington, USA), and the integrity of the RNA was checked on a 1.7% agarose gel (w/v). Aliquots of RNA from each sample were used to create a mixed pooled sample with equal concentrations of extracted RNA.

Before library preparation, the total RNA of the mixed sample was analyzed using the 4200 TapeStation system (Agilent; Santa Clara, USA) for integrity and quantified using a fluorometric assay using the Qubit™ RNA Broad-Range Kit (Invitrogen; Massachusetts, USA). cDNA synthesis, library construction, and sequencing were performed at the Laboratório Central de Tecnologias de Alto Desempenho em Ciências da Vida (LaCTAD) at the Universidade Estadual de Campinas, São Paulo, Brazil. The sequencing library was prepared following the manufacturer’s protocol with reagents supplied by Illumina’s Stranded mRNA Prep, Ligation (Illumina; San Diego, USA) and sequenced using Illumina NextSeq 2000 (Illumina) technology, generating 150 base pair (bp)-long, paired-end reads.

Raw RNA sequence reads were preprocessed to trim adapter sequences using BBMap v39.08 ^43^. Reads with a Phred quality score of ≥30 were retained for de novo transcriptome assembly with the Trinity suite v2.15.2^44^ using parameters optimized for paired-end reads. The open reading frames (ORFs) of the assembled transcripts were predicted using TransDecoder v5.7.1.^45^ ORFs encoding peptides longer than 100 amino acids were screened for homologous proteins in the non-redundant (NR) database using Diamond BLASTp v2.1.9.^46^ Conserved domains were identified with HMMER v3.4 ^47^ by searching against the Pfam-A v2024-05-28 database.^48^ The results from Pfam and NR were used for the second step of the TransDecoder pipeline (TransDecoder.Predict). The final protein description annotations were performed using OmicsBox software v1.0.34.

### 2.5. Protein extraction

For proteomic analysis, three biological replicate samples (each containing 300 mg FM) of seeds collected at 4, 6, and 8 WAA before germination (time 0; 8 WAA_0h) and seeds collected at 8 WAA after 24 h of germination (8 WAA_24h) were obtained. The samples were flash frozen in liquid nitrogen and stored at −80 °C until analysis.

Protein extraction was carried out using the sequential solubility classification method according to Romero-Rodríguez et al.^49^, with some modifications. Briefly, the samples were pulverized in liquid nitrogen using a ceramic mortar and pestle and washed three times with petroleum ether (SigmaCAldrich) to reduce the lipid content. The resulting pellet was then resuspended in a water-soluble protein extraction buffer (FR1) composed of 10 mM Tris-HCl (pH 7.5; Cytiva), 10 mM MgCl_2_.6H_2_O (Synth; São Paulo, Brazil), 10 mM CaCl_2_.2H_2_O (Synth), and 0.1% (w/v) DTT (Cytiva). The extract was continuously stirred for 1 h at 4 °C, followed by centrifugation at 15,000 × g for 15 min at 4 °C (all the stirring and centrifugation steps of protein extraction were performed under these conditions unless otherwise noted), and the supernatant was collected in a new tube. The pellet remaining after the extraction of water-soluble protein extraction was resuspended in saline-soluble protein extraction buffer (FR2), consisting of 10 mM Tris-HCl, 1 M NaCl (Vetec; Rio de Janeiro, Brazil), 10 mM EDTA (Sigma-Aldrich), and 0.1% DTT (w/v; Cytiva). The extract was continuously stirred, followed by centrifugation, and the supernatant was collected in a new tube. The saline extraction pellet was subsequently resuspended in 75% aqueous ethanol solution (Merck; Darmstadt, Germany) (FR3). The extract was stirred continuously for 1 h at 4 °C and then centrifuged at 15,000 × g for 15 min at 4 °C, after which the supernatant containing the ethanol-soluble proteins was transferred to a new tube. Subsequently, the ethanolic extraction pellet was resuspended in alkaline protein extraction buffer (FR4) composed of 0.1 M NaOH (Merck). The extract was continuously stirred for 1 h at 4 °C and then centrifuged at 15,000 × g for 15 min at 4 °C, and the resulting supernatant was transferred to a new tube. Finally, we enhanced our workflow with an additional sequential step to recover total residual proteins, including membrane- associated proteins and other proteins that were not soluble in previous buffers. The alkaline extraction pellet was resuspended in chaotropic protein extraction buffer (FR5), consisting of 7 M urea (Cytiva), 2 M thiourea (Cytiva), 1% DTT (Cytiva), 2% Triton-100 (Cytiva), and 1 mM PMSF (SigmaCAldrich). The extract was continuously stirred and then centrifuged at 15,000 × g for 15 min at 4 °C, after which the supernatant was transferred to a new tube. The protein content of all fractions (FR1CFR5) was determined by using a 2-D Quant Kit (Cytiva). A scheme of the third extraction methodology is shown in Figure 1.

**Fig. 1.**
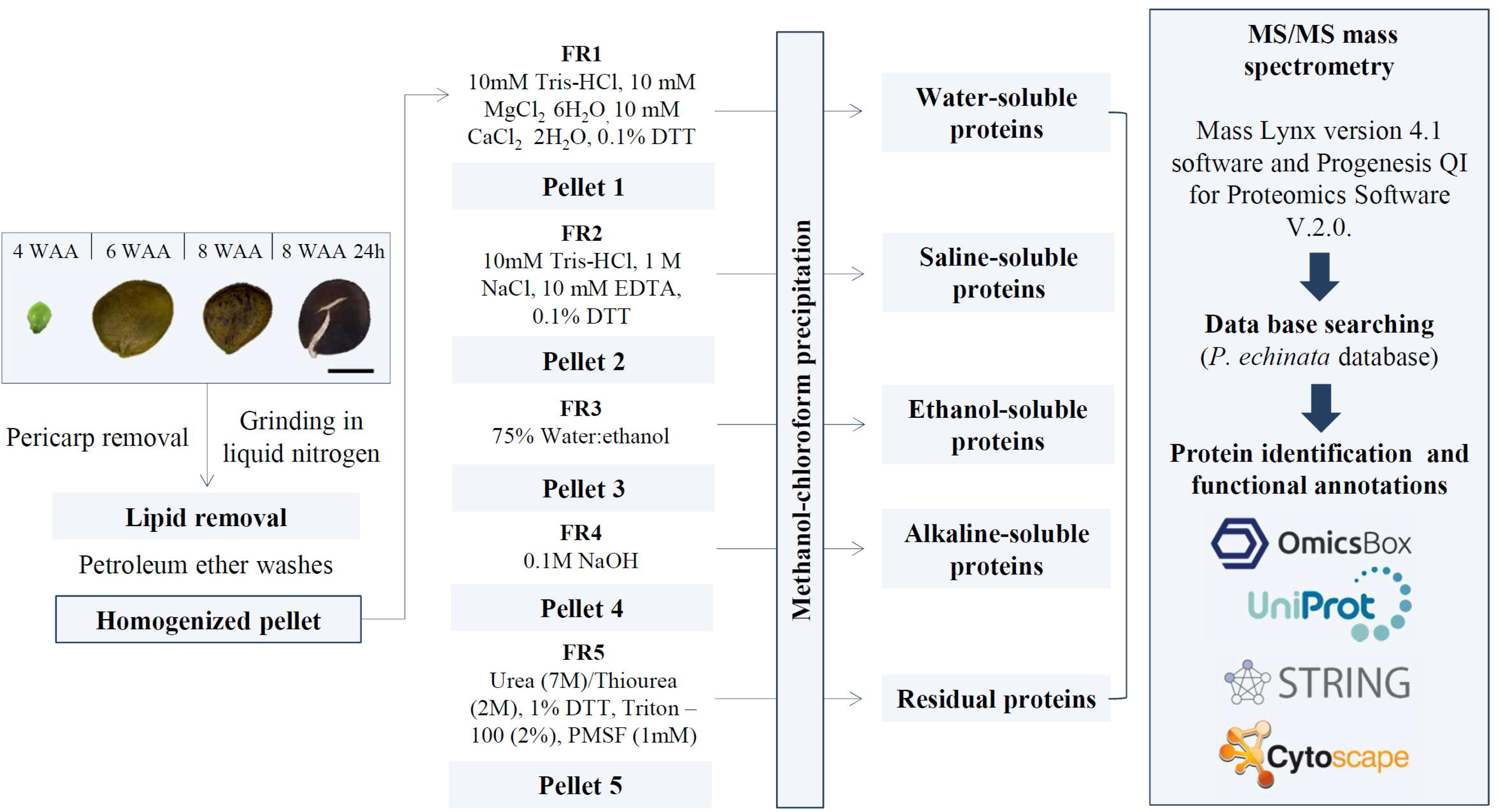
Flowchart of the protein extraction procedure for *P. echinata* seeds and MS/MS analysis. FR1 = water-soluble proteins; FR2 = saline-soluble proteins; FR3 = ethanol-soluble proteins; FR4 = alkaline- soluble proteins; FR5 = residual proteins.

### 2.6. Protein Digestion and Proteomic Analysis

All fractions of each biological replicate were individually digested after carefully indexed (identified) for subsequent processing of the samples as a fractionated experiment. First, protein fractions were precipitated using the methanol/chloroform method.^50^ After protein precipitation, the samples were resuspended in 7 M urea/2 M thiourea buffer, and then trypsin digestion (100 ng µL^-1^; V5111; Promega; Madison, USA) was performed using filter-aided sample preparation (FASP) methodology.^51^ The peptides were subsequently resuspended in 30 µL of a solution containing 95% water (Tedia; Fairfield, USA), 5% acetonitrile (Sigma-Aldrich), and 0.1% formic acid (Sigma–Aldrich) and quantified using the A205 nm protein and peptide methodology using a NanoDrop 2000c spectrophotometer (Thermo Fisher Scientific; Waltham, USA). The prepared fraction samples were transferred to Total Recovery Vials (Waters; Manchester, UK) for mass spectrometry analysis.

Mass spectrometry analysis was performed using a nanoAcquity UPLC system connected to a Q- TOF SYNAPT G2-Si instrument (Waters). The runs involved the injection of 1 µg of peptide from each fraction into all three biological replicates. During separation, the samples were loaded into a nanoAcquity UPLC M-Class Symmetry C18 5 μm trap column (180 μm × 20 mm; Waters) at 5 µL min^-1^ for 3 min and then onto a nanoAcquity M-Class HSS T3 1.8 μm analytical reversed-phase column (75 μm × 150 mm; Waters) at 400 nL min^-1^, with a column temperature of 45 °C. For the peptide elution, a binary gradient was used, with mobile phase A consisting of water and 0.1% formic acid (Sigma-Aldrich) and mobile phase B consisting of acetonitrile (Sigma-Aldrich) and 0.1% formic acid (Sigma-Aldrich). The gradient elution started at 5% B, increased from 5 to 41% B until 92 min, increased again from 41 to 97% B until 96 min, remained at 97% B until 100 min, decreased to 5% B until 102 min, and finally remained at 5% B until the end of the run, at 118 min. Mass spectrometry was performed in positive and resolution mode (V mode), with 35,000 full widths at half maximum and ion mobility separation (IMS), and in data- independent acquisition mode (HDMS^E^). The ion mobility wave was set to a velocity of 650 m s^-1^, and the helium and IMS gas flows were 180 and 90 mL min^-1^, respectively. The transfer collision energy increased from 19 to 55 V in high-energy mode; the cone and capillary voltages were 40 and 2800 V, respectively; and the source temperature was 80 °C. For the time-of-flight (TOF) parameters, the scan time was set to 0.5 s in continuum mode with a mass range of 50-2000 Da. Human [Glu^1^]-fibrinopeptide B at 100 fmol µL^-1^ was used as an external calibrant, and lock mass acquisition was performed every 30 s. Mass spectra were acquired by Mass Lynx v4.1 software.

Spectral processing and database search were performed using the fractionation analysis workflow with Progenesis QI for Proteomics Software v2.0 (Nonlinear Dynamics; Newcastle, UK). To analyze all fraction data, they were organized in a combined analyzed fraction experiment. This was followed by the analysis of each single fraction and then recombining the analyzed fractions into a multifraction experiment. The processing parameters were set to a low energy threshold of 150 (counts), an elevated energy threshold of 50, an intensity threshold of 750, two missed cleavages, a minimum fragment ion per peptide equal to three, a minimum fragment ion per protein equal to seven, a minimum peptide per protein equal to two, fixed modifications of carbamidomethyl and variable modifications of oxidation and phosphoryl (STY). The false discovery rate (FDR) for peptide and protein identification was set to a maximum of 1%, with a minimum peptide length of six amino acids and a maximum mass error of 10 ppm. For protein identification, two distinct databases were used: a) the non-redundant *P. echinata* database generated from RNA-seq data and b) the *Glycine max* (ID: UP000008827) protein sequence database from UniProtKB (http://www.uniprot.org). BLAST analysis was performed to compare identified proteins with *Glycine max* and identified proteins with the *Paubrasilia echinata* databank using the NCBI BLAST+ suite^52^ within the R software environment v4.4.1.^53^ The mass spectrometry proteomics data and the non-redundant protein database of *P. echinata,* generated from RNA sequence data, have been deposited in the ProteomeXchange Consortium^54^ via the PRIDE^55^ partner repository with the dataset identifier PXD056397.

Label-free quantification was estimated using the TOP3 quantification approach. To ensure the quality of the results after data processing, only proteins that were consistently present or absent (for unique proteins) in all three biological replicates were considered for differential accumulation analysis. Data were analyzed using Student’s t test (two-tailed). Proteins with a significant t-test (P<0.05) were considered up-accumulated if the log2 value of the fold change (FC) was greater than 0.60 and down-accumulated if the log2 value of the FC was less than −0.60. Functional annotations were performed using OmicsBox software v1.0.34 and UniProtKB (http://www.uniprot.org). For the development of the seeds, the Mfuzz package^56^ was used for clustering analysis of DAPs in at least one comparison between the harvested times studied. Functional enrichment analysis of the DAPs was performed using the *Arabidopsis thaliana* reference database, employing a STRING search^57^ with confidenceC>C0.7 and Cytoscape v3.10.2.^58^

### 2.7. Free polyamine (PA) analysis

For the analysis of PA, three biological replicate samples (each containing 200 mg FM) of seeds collected at 4, 6, and 8 WAA before (time 0) and at 6, 12, 24, and 96 h after germination were used. These samples were flash frozen in liquid nitrogen and stored at -20 °C until analysis. Free PAs were determined according to the methods of Aragão et al.^59^. The samples were macerated in 1.2 mL of 5% perchloric acid (Merck). After 1 h of incubation at 4 °C, the samples were then centrifuged for 20 min at 20,000 × g at 4 °C. The supernatant was collected, and the free PAs were analyzed by dansylation with dansyl chloride (Merck). Subsequently, PAs were identified and quantified by high-performance liquid chromatography (HPLC; Shimadzu) using a 5 μm C18 reversed-phase column (Shin-pack CLC ODS; Shimadzu).

The gradient for the mobile phase was generated by progressively adding higher volumes of absolute acetonitrile (Merck) to a 10% aqueous solution of acetonitrile (pH 3.5 adjusted with HCl). The gradient of absolute acetonitrile was programmed to 65% in the first 10 min, increasing from 65 to 100% between 10 and 13 min and maintaining 100% between 13 and 21 min at a flow rate of 1 mL min^-1^ at 40 °C. PA peaks were detected using a fluorescence detector set at wavelengths of 340 nm (excitation) and 510 nm (emission). Peak areas and PA retention times were measured by comparison with standard Put, Spd, and Spm PAs (Sigma-Aldrich). Additionally, 0.05 mM 1,7-diaminoheptane (DAH) (SigmaCAldrich) was used as an internal standard.

### 2.8. Data Analysis

All experiments were carried out according to a completely randomized design. For data analysis, the assumptions of normality and homogeneity of the treatment variances were first verified by the Shapiro-Wilk test and the Bartlett test, respectively. The data was then subjected to analysis of variance (ANOVA), and the means were separated by Tukey’s test (P<0.05). The parameters were analyzed with the aid of the R software environment v. 4.3.0 (R Core Team, 2023) with the aid of the easyanova package.^60^

## 3. Results

### 3.1. Seed development and germination

We observed a distinct length, fresh matter (FM), and dry matter (DM) of *P. echinata* seeds over time (Figure 2A). For the three measured parameters analyzed, there was a significant increase from 4 to 6 WAA, followed by a slight decrease to 8 WAA (Figure 2B-D).

**Fig. 2.**
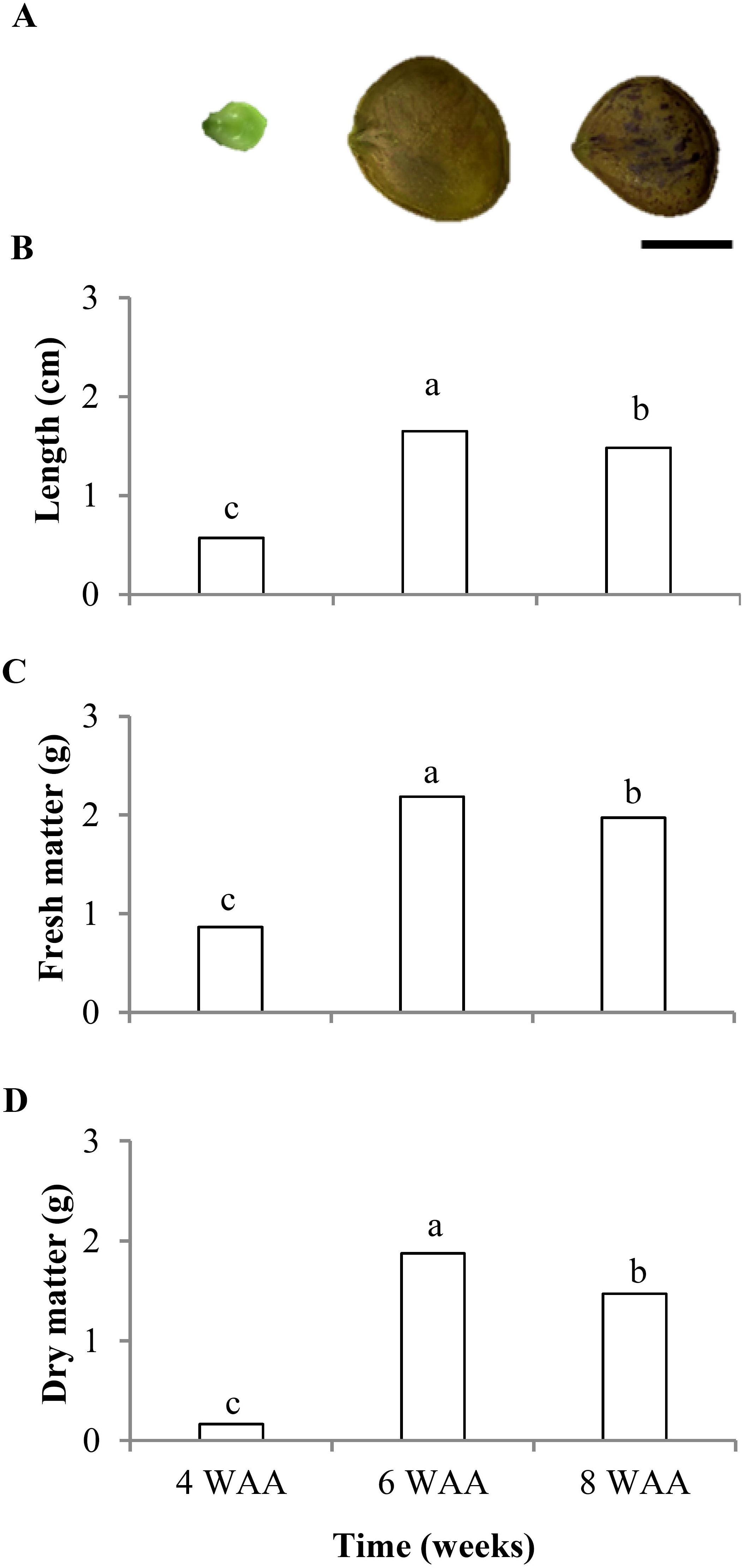
Morphological aspects (A), length (B), fresh matter (C), and dry matter (D) of *P. echinata* immature seeds at 4 and 6 weeks after anthesis (WAA) and mature dispersed seeds at 8 WAA. Means followed by different letters denote significant differences according to Tukey’s test (P < 0.05). CV = Coefficient of variation. (n = 5; CV of length = 3.54%; CV of FM = 4.75%; CV of DM = 22.62%). Bar A = 1 cm.

*P. echinata* seeds have a short imbibition phase and emit radicles within the first 24 h, with the formation of well-developed seedlings after eight days of germination (192 h; Figure 3A). Seeds harvested at 8 WAA had a higher percentage of germination and GSI than seeds harvested at 6 WAA, while seeds harvested at 4 WAA did not germinate (Figure 3B-C).

**Fig. 3.**
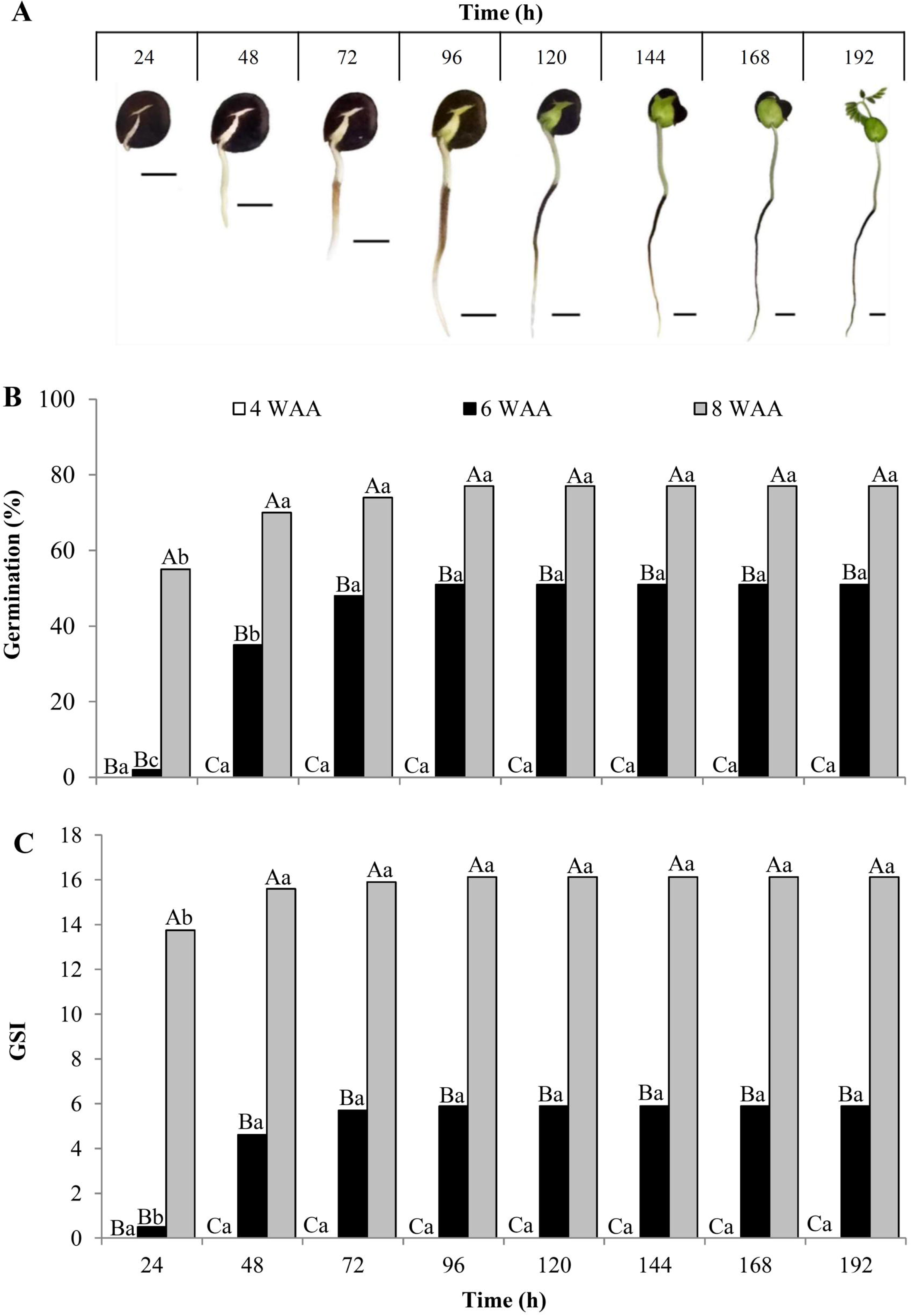
Morphological characteristics of *P. echinata* seeds at 8 weeks after anthesis (WAA) and of seedlings at 192 h after germination (A), germination percentage (B), and germination speed index (C) of immature seeds at 4, 6, and 8 WAA. The uppercase letters indicate significant differences between the seeds harvested from different WAAs at the same time (h) of germination. Lowercase letters indicate significant differences between the different times of germination (h) for seeds harvested at each WAA. Means followed by different letters indicate significant differences according to Tukey’s test (P < 0.05). CV = coefficient of variation (n = 4; CV of germination = 17.66%; CV of GSI = 15.15%). Bars in A= 1 cm.

### 3.2. Development of a non-redundant protein database of *P. echinata*

A total of 57,212,134 high-quality RNA-Seq reads were utilized for transcriptome assembly, resulting in 83,040 transcripts. The completeness analysis performed with BUSCO indicated that 82.7% of the transcripts were complete. TransDecoder prediction identified 55,426 protein sequences (PRIDE identifier PXD056397).

For the proteomic analysis of seed development, the database generated for *P. echinata* presented a significant increase in the number of total identified proteins, with 2250 proteins (Table S1), compared to the 1076 proteins identified through the *G. max* database (Table S2). For comparative proteomics between seeds 8 WAA after 24 h of germination and those 8 WAA, 2244 proteins were identified via the *P. echinata* database (Table S3), while 1077 proteins were detected via the *G. max* database (Table S4).

BLAST analysis revealed that the proteins identified in *P. brasilia* database covered most of the proteins identified from *G. max*, with a high percentage of shared identity in both proteomic analyses (Tables S5- S6).

On the basis of these results, all comparative proteomic analyses were performed using the protein database generated for *P. echinata*.

### 3.3. Protein profile during seed development

The sequential extraction method showed variations in protein concentrations from the different extraction fractions and harvest times (Figure 4). At 4 WAA, the seeds consisted predominantly of alkaline-soluble proteins (FR4) and residual proteins (FR5). On the contrary, seeds harvested at 6 and 8 WAA presented a relatively high content of water-soluble proteins (FR1), with substantial amounts of soluble saline (FR2), soluble alkaline (FR4), and residual (FR5) proteins. The ethanol-soluble proteins (FR3) did not differ significantly at any point during harvest.

**Fig. 4.**
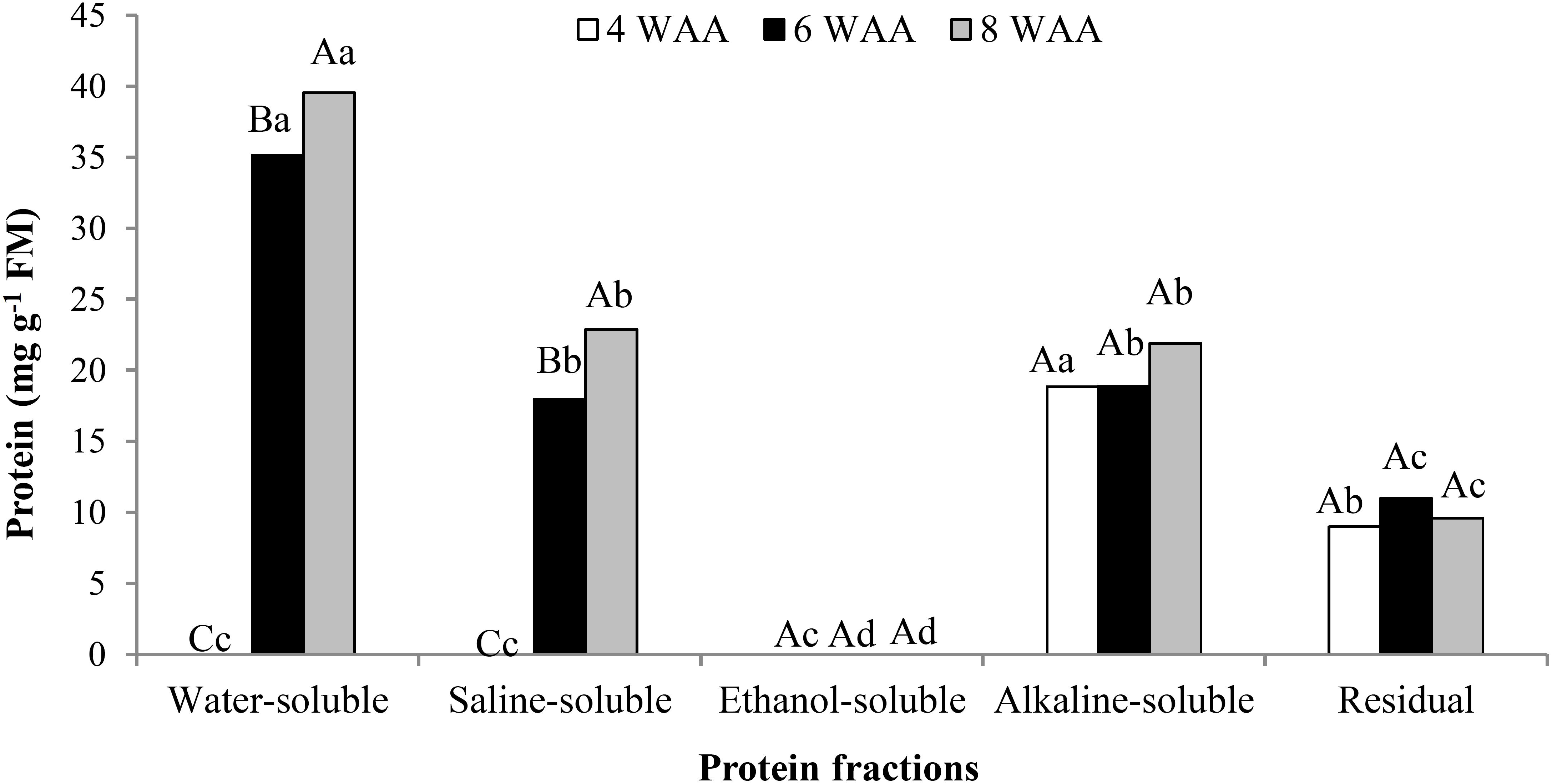
Protein quantification (mg g^-1^ FM) in *P. echinata* seeds harvested at 4, 6, and 8 weeks after anthesis (WAA) across different extraction fractions. The uppercase letters indicate significant differences between seeds harvested at different WAA within the same extraction fraction. Lowercase letters indicate significant differences among extraction fractions for seeds harvested at the same WAA. Different letters denote significant differences according to Tukey’s test (P < 0.05). CV = coefficient of variation. (n = 3; CV = 15.38%).

Of the 2250 proteins identified in seeds harvested in 4, 6, and 8 WAA, 1104 proteins were differentially accumulated (DAPs) in at least one of the comparisons (6 WAA/4 WAA or 8 WAA/6 WAA) (Table S1). Cluster analysis of DAPs revealed four distinct clusters representing unique protein accumulation patterns during seed development (Figure 5; Table S1). Cluster 1 comprises 168 proteins whose accumulation increased at all analyzed time points. Cluster 2 comprises 703 proteins whose abundance increased between 4 and 6 WAA, followed by stabilization from 6 to 8 WAA. Cluster 3 comprises 96 proteins that were most abundant at 4 WAA, their abundance declining at 6 WAA and then stabilizing at 8 WAA. Cluster 4 comprises 52 proteins with an irregular accumulation pattern, where protein levels decreased from 4 to 6 WAA and subsequently increased from 6 to 8 WAA.

**Fig. 5.**
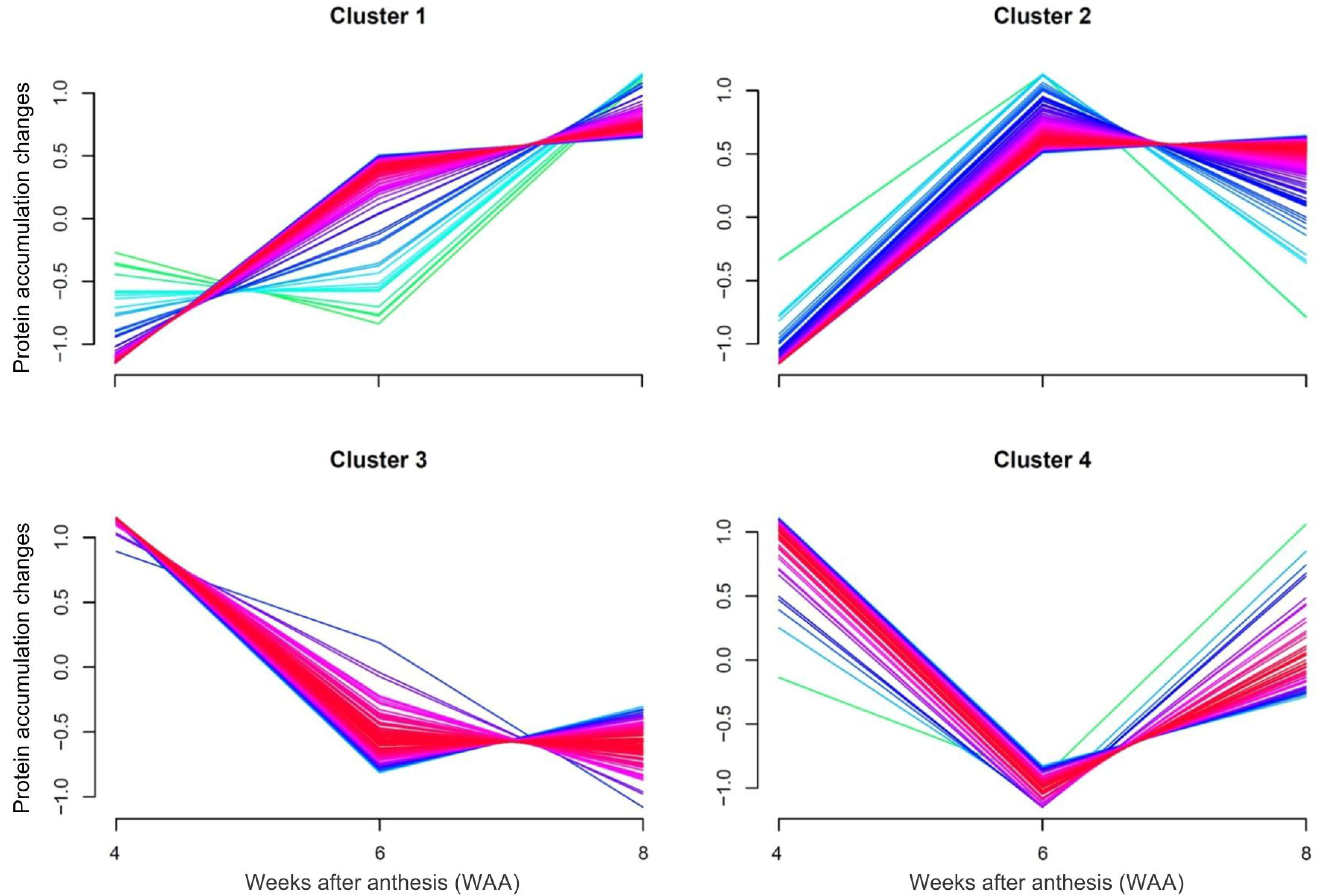
Comparative analysis of proteomics data during the development of *P. echinata* seeds, revealing four distinct protein accumulation patterns: proteins showing an increase in accumulation across all analyzed time points (Cluster 1); proteins with a marked increase between 4 and 6 weeks after anthesis (WAA), followed by stabilization from 6 to 8 WAA (Cluster 2); proteins most abundant at 4 WAA, with abundance declining at 6 WAA and then stabilizing at 8 WAA (Cluster 3); and proteins exhibiting an irregular pattern of accumulation, characterized by a decrease in accumulation from 4 to 6 WAA, followed by an increase from 6 to 8 WAA (Cluster 4).

For Cluster 1, the enrichment analysis highlighted 88 biological processes (Table S7) and the PPI network (Figure 6A) revealed consistent processes such as response to stress (51 proteins), protein metabolic process (42 proteins), cellular catabolic process (22 proteins), translation (20 proteins), and carbohydrate derivative metabolic process (14 proteins). Cluster 2 was characterized by a broader range of enriched processes, resulting in 196 biological processes (Table S7). Cluster 2 PPI networks (Figure 6B) were associated with the cellular metabolic process (329 proteins), the response to stress (170 proteins), the protein metabolic process (147 proteins), the carboxylic acid metabolic process (95 proteins), the translation (76 proteins), and the carbohydrate derivative metabolic process (53 proteins). For Cluster 3, the enrichment analysis identified 20 biological processes (Table S7) and the PPI network (Figure 6C) highlighted key processes, including photosynthesis (14 proteins), the carbohydrate derivative biosynthetic process (9 proteins), and flavonoid biosynthetic process (7 proteins). For Cluster 4, the enrichment analysis identified 10 biological processes (Table S7); however, no enriched PPI network was observed via Cytoscape.

**Fig. 6.**
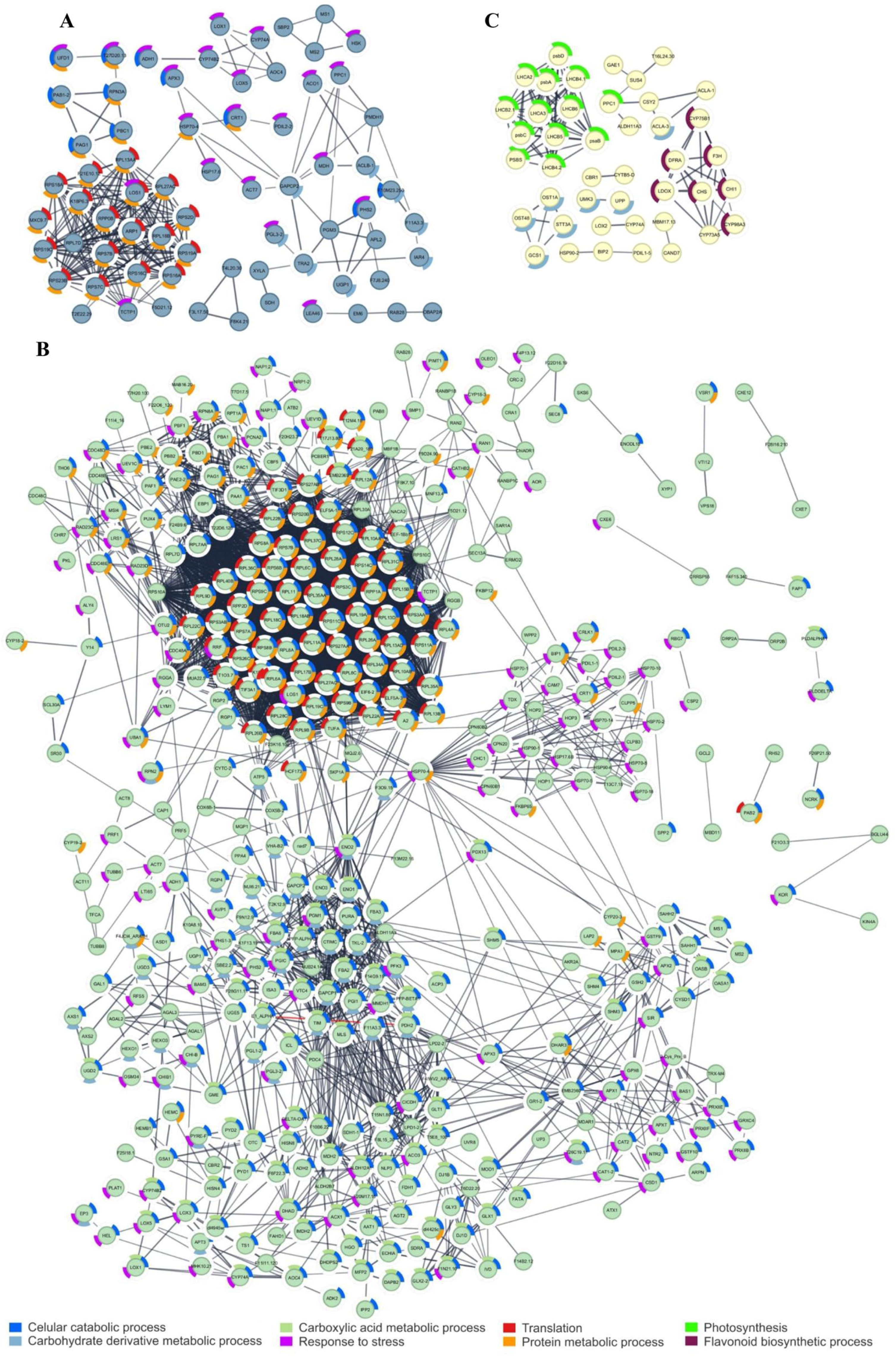
Functional analysis of differentially accumulated proteins (DAPs) during seed development of *P. echinata*, conducted via STRING within Cytoscape, highlighting the main biological processes enriched for Cluster 1—proteins with consistently increasing accumulation (A), Cluster 2—proteins with a marked increase between 4 and 6 weeks after anthesis (WAA), followed by stabilization (B), and Cluster 3— proteins most abundant at 4 WAA, with a subsequent decline and stabilization (C). No biological process enrichment was observed for Cluster 4.

### 3.4. Differential protein accumulation during seed development

The development of *P. echinata* seeds from 4 to 8 WAA is regulated by the differential accumulation of proteins. Among them, the proteins of the eukaryotic translation initiation factor 3 subunit A (eIF3A; paubrasilia_39643) and eIF5A (paubrasilia_33473) were up-accumulated at 6 compared to 4 WAA. Additionally, the proteins eIF1A (paubrasilia_08033), eIF3D (paubrasilia_30254), and eIF6 (paubrasilia_37410) were uniquely detected in seeds at 6 WAA. Furthermore, 18 DAPs associated with proteasome activity were identified in the 6 WAA/4 WAA comparison. Among these, the E3 ubiquitin-protein ligase (paubrasilia_15904), proteasome subunit alpha (paubrasilia_42692, paubrasilia_46179, paubrasilia_49676, and paubrasilia_55364), and proteasome subunit beta (paubrasilia_45947 and paubrasilia_51037) genes were unique to the seeds at 6 WAA. On the contrary, the ubiquitin-activating enzymes E1 (paubrasilia_53937 and paubrasilia_53942), the E3 ubiquitin-protein ligase (paubrasilia_04063), the proteasome subunits alpha (paubrasilia_19415, paubrasilia_42693, and paubrasilia_18688), and the proteasome subunits beta (paubrasilia_37204, paubrasilia_04966, paubrasilia_24225, and paubrasilia_32732) were up-accumulated in seeds at 6 compared to those at 4 WAA.

Additionally, seven homologs of cell division cycle protein 48 (CDC48) were differentially regulated in the 6 WAA/4 WAA comparison. Of these, six were unique to 6 WAA (paubrasilia_43841, paubrasilia_21736, paubrasilia_47378, paubrasilia_50791, paubrasilia_50804, and paubrasilia_50808), and one was up-regulated (paubrasilia_50794).

Some enzymes associated with the TCA cycle, such as aconitate hydratase (paubrasilia_32357 and paubrasilia_16147), malate synthase (paubrasilia_26382), malate dehydrogenase (paubrasilia_26140, paubrasilia_17815, paubrasilia_26382 and paubrasilia_36465), succinate-CoA ligase (paubrasilia_41056 and paubrasilia_41050), and succinate dehydrogenase (paubrasilia_39985), were up-accumulated in seeds at 6 WAA/4 WAA comparison, while isocitrate lyase (paubrasilia_16732) and succinate-CoA ligase (paubrasilia_37976) were unique to seeds at 6 WAA.

In *P. echinata*, key glycolytic proteins such as phosphoglycerate kinase (paubrasilia_13254), enolase (paubrasilia_13506, paubrasilia_25590, paubrasilia_25591, paubrasilia_41363), glyceraldehyde-3- phosphate dehydrogenase (paubrasilia_19718, paubrasilia_19720, paubrasilia_15333, paubrasilia_06794, paubrasilia_06795), triosephosphate isomerase (paubrasilia_02133, paubrasilia_02134 paubrasilia_55147), fructose-bisphosphate aldolase (paubrasilia_15563, paubrasilia_47698, paubrasilia_28173), glucose-6-phosphate isomerase (paubrasilia_32316, paubrasilia_34650), and glucose- 6-phosphate 1-epimerase (paubrasilia_29237, paubrasilia_29368) were either up-accumulated or unique in seeds at 6 compared with those at 4 WAA, suggesting that these proteins may also play a role in seed desiccation.

Additionally, 15 LEA proteins were found to be DAPs. Furthermore, these proteins were not specifically enriched in any particular biological process, 14 of which were either up-accumulated or unique to seeds at 6 compared to those at 4 WAA (Clusters 1 and 2). Additionally, six of these proteins (paubrasilia_03731, paubrasilia_24605, paubrasilia_30609, paubrasilia_51099, paubrasilia_51372, and paubrasilia_51374) were up-accumulated in both comparisons: 6 WAA/4 WAA and 8 WAA/6 WAA. Proteins related to protein folding and assembly were up-accumulated or unique to seeds collected at 6 WAA compared to those collected at 4 WAA (Cluster 2). Among these proteins, four are disulfide- isomerase proteins, and 22 are heat shock proteins (HSPs).

Proteins associated with photosystem I (paubrasilia_14331, paubrasilia_06896, and paubrasilia_15968) and photosystem II (paubrasilia_24320, paubrasilia_00600, paubrasilia_14238, and paubrasilia_00599) were down-accumulated in seeds collected at 6 compared to those collected at 4 WAA (Cluster 3). Furthermore, key enzymes involved in the flavonoid biosynthetic pathway, including chalcone isomerase (paubrasilia_04859), chalcone synthase 1-like (paubrasilia_42058), dihydroflavonol- 4-reductase (paubrasilia_48528 and paubrasilia_37268), leucoanthocyanidin dioxygenase (paubrasilia_26029), and flavonoid 3’-monooxygenase-like (paubrasilia_45495), were down-accumulated in the 6 WAA/4 WAA comparison.

Furthermore, four proteins up-accumulated in the 6 WAA/4 WAA comparison from Cluster 2 were associated with methionine synthesis, a precursor of PA synthesis, with two adenosylhomocysteinases (paubrasilia_07653 and paubrasilia_07655) and two 5-methyltetrahydropteroyltriglutamateChomocysteine methyltransferases (paubrasilia_31470 and paubrasilia_41057).

### 3.5. Protein profile during seed germination

During germination, water-soluble proteins (FR1), soluble saline (FR2), soluble alkaline (FR4), and residual (FR5) proteins were successfully identified, but none of the ethanol-soluble proteins (FR3 fraction) was detected (Figure 7). Before starting germination (8 WAA_0h), the protein content in these four fractions was higher than that of the seeds at 24 h of germination, after root protrusion (Figure 7). The FR1 fraction had a significantly higher protein concentration before and after germination, highlighting the potential role of water-soluble proteins in the early stages of the development of *P. echinata* seedlings.

**Fig. 7.**
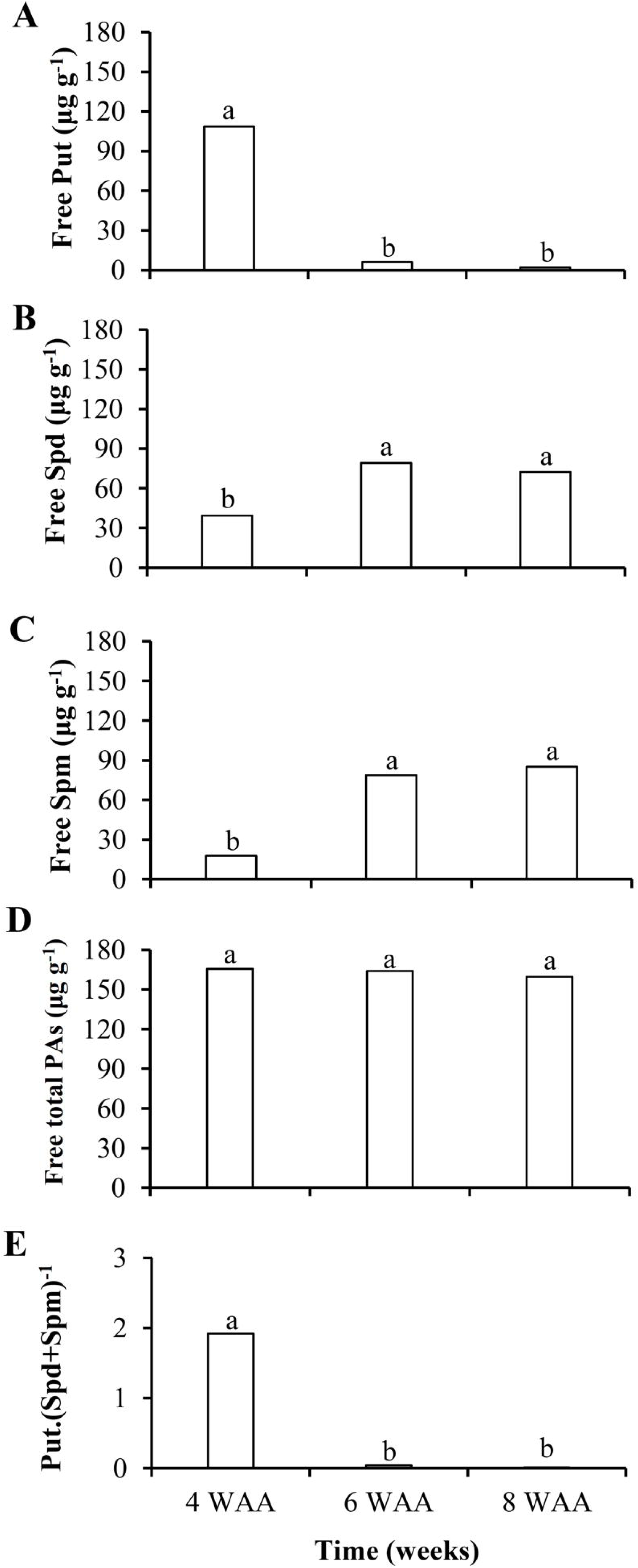
Protein contents (mg g^-1^ FM) of *P. echinata* seeds harvested at 8 weeks after anthesis (WAA) before (time 0; 8 WAA) and after 24 h of germination (8 WAA_24h) in different extraction fractions. The uppercase letters indicate significant differences at different germination times (0 and 24 h) in the same extraction fraction. Lowercase letters indicate significant differences across the different extraction fractions at the same time of germination (0 or 24 h). Different letters indicate significant differences according to Tukey’s test (P < 0.05). CV = coefficient of variation (n = 3; CV = 11.19%).

Of the 2244 proteins identified in early seed germination, 145 were DAPs in seeds at 8 WAA_24h/8 WAA_0h. Among the DAPs, 52 were up-accumulated, and 93 were down-accumulated (Table S3). The enrichment analysis of the DAPs revealed 28 biological processes (Table S8), while the PPI network emphasized key processes such as the carboxylic acid metabolic process (11 proteins), the organic substance biosynthetic process (21 proteins), protein folding (6 proteins), response to stress (20 proteins) and the small molecule metabolic process (16 proteins) (Figure 8).

**Fig. 8.**
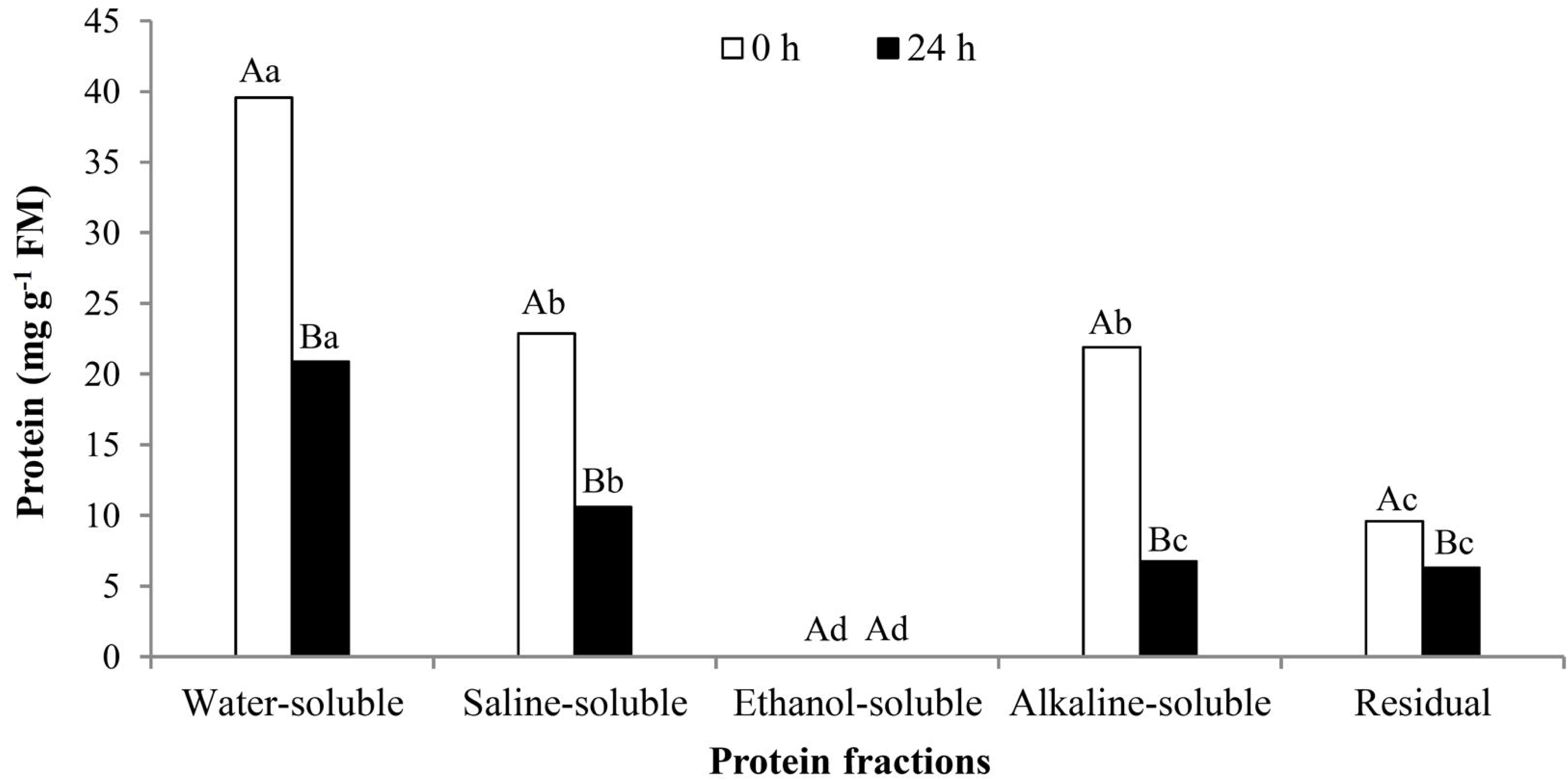
Functional analysis of differentially accumulated proteins (DAPs) during seed germination in *P. echinata* conducted via STRING within Cytoscape.

### 3.6. Differential protein accumulation during seed germination

Proteomic analysis revealed a higher number of down-accumulated proteins than up-accumulated at 8 WAA_24h after imbibition than at 8 WAA_0h before imbibition. Among them, 4 LEA proteins (paubrasilia_03973, paubrasilia_28787, paubrasilia_30609, and paubrasilia_03731) were down-accumulated in seeds at 24 h of imbibition compared to 8 WAA seeds before incubation. Moreover, protein folding and response to stress, two of the identified enriched biological processes, revealed that the majority of proteins were down-accumulated in the seeds at 24 h post-imbibition compared to before incubation (8 WAA_24h/8 WAA_0h comparison), including two HSPs (paubrasilia_14874 and paubrasilia_47886).

Furthermore, three other enriched processes, the small molecule metabolic process, the carboxylic acid metabolic process, and the organic substance biosynthetic process, showed that most proteins were down-accumulated in *P. echinata* seeds during germination (8 WAA_24h/8 WAA_0h comparison). Among them, allene oxide cyclase 4 (paubrasilia_14955), two 5-methyltetrahydropteroyltriglutamate-homocysteine S-methyltransferases (paubrasilia_41067 and paubrasilia_41060), and the SAM-binding methyltransferase (paubrasilia_49529) enzymes were down-accumulated in the seeds at 24 h after imbibition compared to before incubation.

### 3.7. PA profile during seed development and germination

The Put content was higher in the seeds harvested at 4 WAA (Figure 9A), while the Spd and Spm contents were higher in the seeds harvested at 6 and 8 WAA (Figure 9B-C). The total free PA content did not vary between the three harvested points of seed development (Figure 9D). The PA ratio Put/(Spd + Spm) was higher in the seeds harvested at 4 WAA (Figure 9E).

**Fig. 9.**
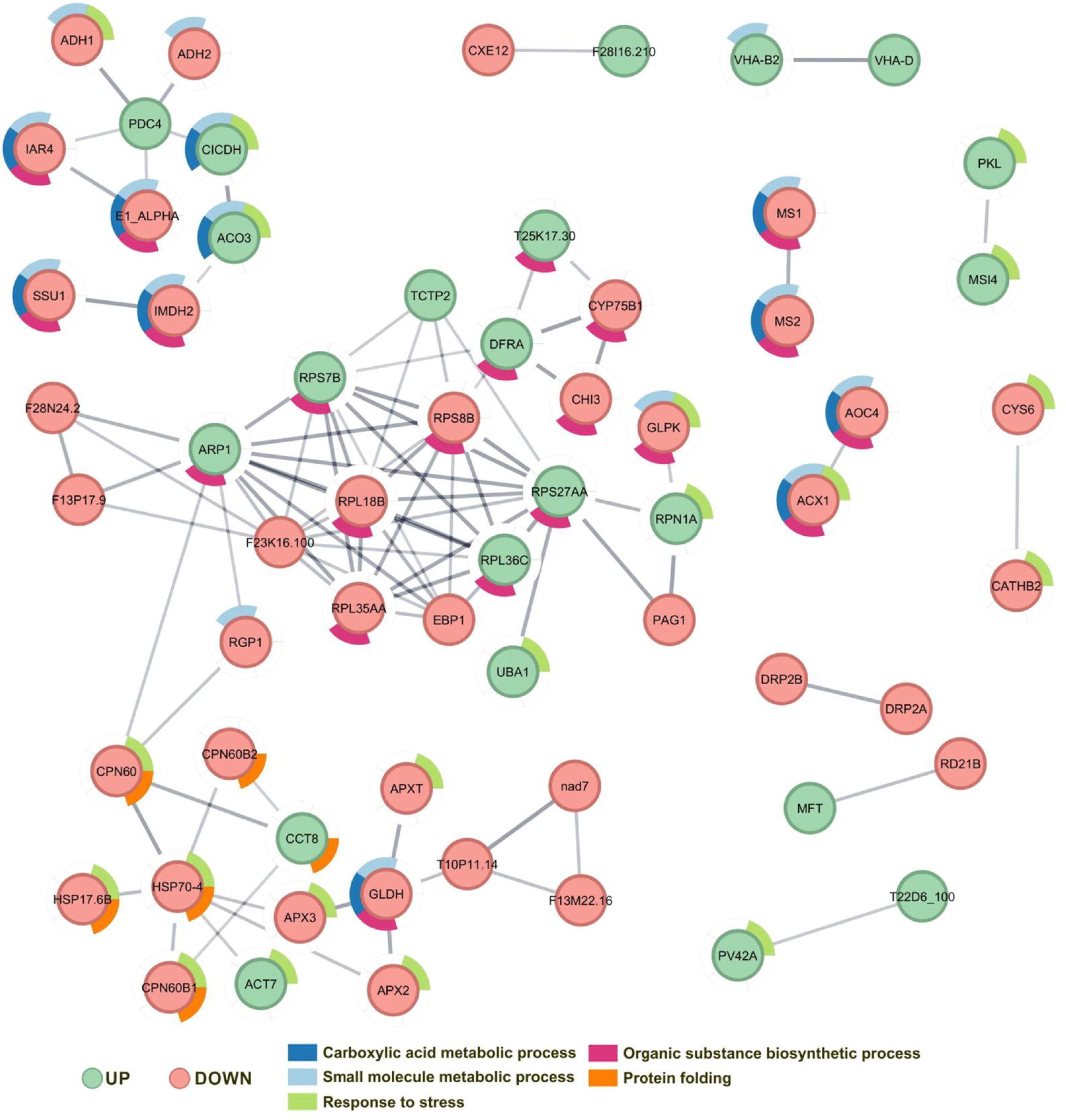
Contents (μg g^-1^ FM) of free Put (A), Spd (B), Spm (C), total free PAs (D), and the PA ratio [ Put.(Spd + Spm)^-1^] (E) in immature *P. echinata* seeds at 4, 6, and 8 weeks after anthesis (WAA). Means followed by different letters indicate significant differences according to Tukey’s test (P < 0.05). CV = coefficient of variation (n = 3; CV of Put = 13.76%, CV of Spd = 12.29%, CV of Spm = 12.64%, CV of total free PAs = 7.92%, CV of PAs ratio = 28.2%).

Among PAs, the Put content was higher in the seeds harvested at 4 WAA at time 0 (before incubation) and decreased after 6 h of incubation, reaching a trend similar to that of the other seeds after 96 h of incubation (Figure 10A). Both the contents of Spd (Figure 10B) and Spm (Figure 10C) were higher in the seeds harvested at 6 and 8 WAA before germination initiation (time incubation). However, these contents decreased after 6 h, reaching lower levels at 96 h of incubation, while the free Spd (Figure 10B) and Spm (Figure 10C) contents in the seeds harvested at 4 WAA remained constant throughout the 96 h germination period.

**Fig. 10.**
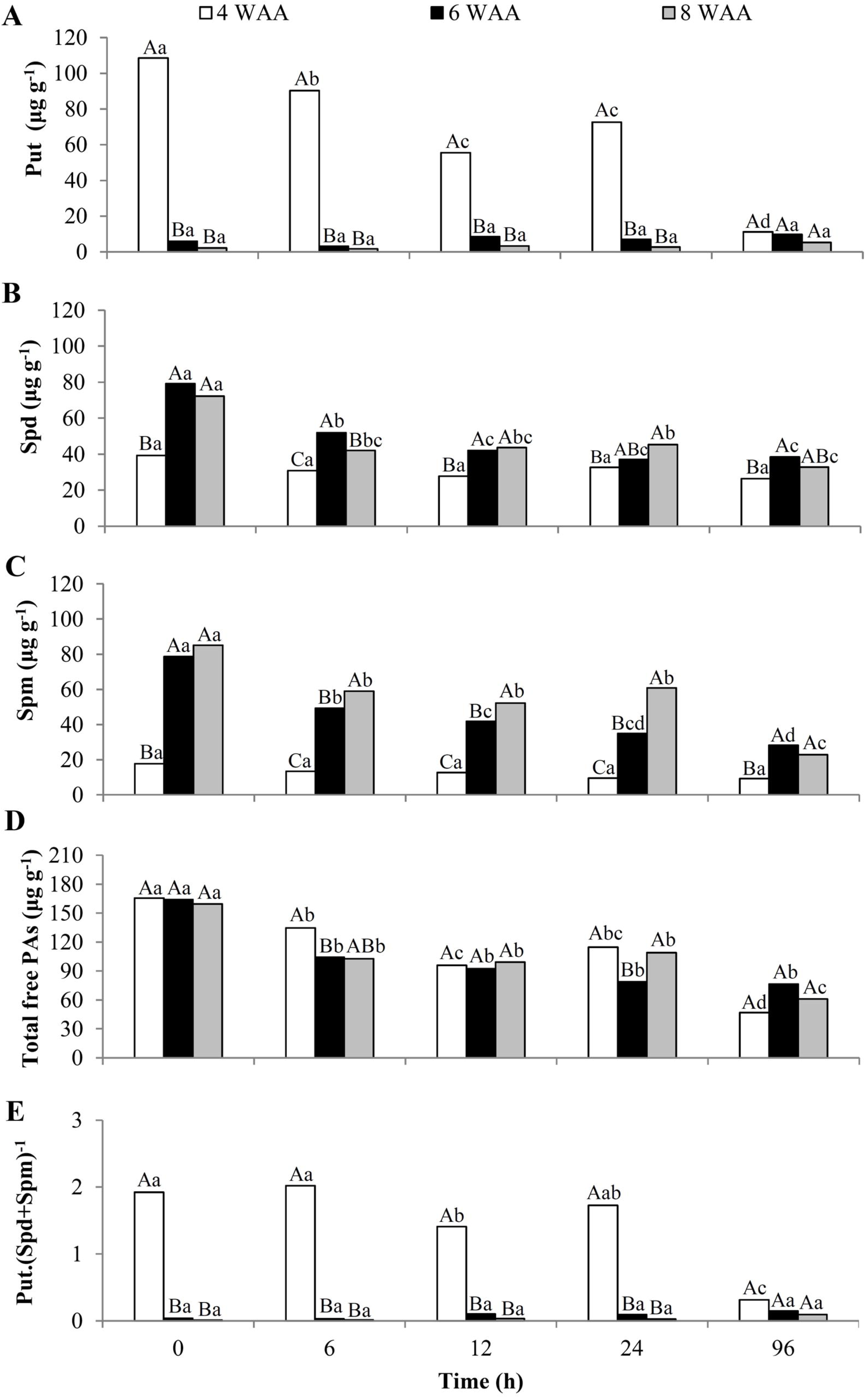
Contents (μg g^-1^ FM) of free Put (A), Spd (B), Spm (C), total free PAs (D), and the PA ratio [Put. (Spd + Spm)^-1^] (E) in P. echinata seeds at 4, 6, and 8 weeks after anthesis (WAA) before (time 0) and after 6, 12, 24, and 96 h of germination. The uppercase letters indicate significant differences between the seeds harvested from different WAAs at the same time (0, 6, 12, 24, or 96 h) of germination. Lowercase letters indicate significant differences between the different germination times (0, 6, 12, 24, and 96 h) for seeds harvested at each WAA. Means followed by different letters denote significant differences according to Tukey’s test (P < 0.05). FM = fresh matter. CV = coefficient of variation. (n = 3; CV of Put = 40.71%; CV of Spd = 13.18%; CV of Spm = 14.14%; CV of free total PAs = 15.44%; CV of PAs ratio = 30.17%).

Total free PA decreased in the seeds at all harvest times (4, 6, and 8 WAA) from 6 to 96 h after germination (Figure 10D). The PA ratio [Put/(Spd + Spm)] was greater in seeds at 4 WAA than in those at 6 and 8 WAA due to their higher Put content. The PA ratio did not differ between the seeds harvested at 6 and 8 WAA (Figure 10E).

## 4. Discussion

### 4.1. Advancement of proteomic analyzes in *P. echinata* through RNA-seq-derived non-redundant protein databases

The protein database generated from RNA-seq of a single pooled sample - comprising different tissues and conditions - represents a powerful strategy to enhance proteomic analyses, ensuring greater coverage and accuracy in protein identification. This study also facilitates a more comprehensive exploration of the identified proteins and the biological processes involved in the development and germination of seeds in *P. echinata*.

These results confirm that non-redundant protein databases derived from RNA-seq data provide a cost-effective and time-efficient alternative to improve protein identification in non-model species lacking a sequenced genome,^22^ as also demonstrated in other species.^61^

### 4.2. Seed development in *P. echinata* is regulated by differential protein accumulation and polyamine dynamics

During development, seeds increase in size as a result of cell proliferation and the accumulation of cellular components on the embryonic axis and storage tissues. However, a reduction in seed size can occur in the final stage of development due to the desiccation process.^62^ This pattern is evident in *P. echinata*, where the length, FM, and DM of mature seeds at 8 WAA are smaller than those at 6 WAA (Figure 11A).

**Fig. 11.**
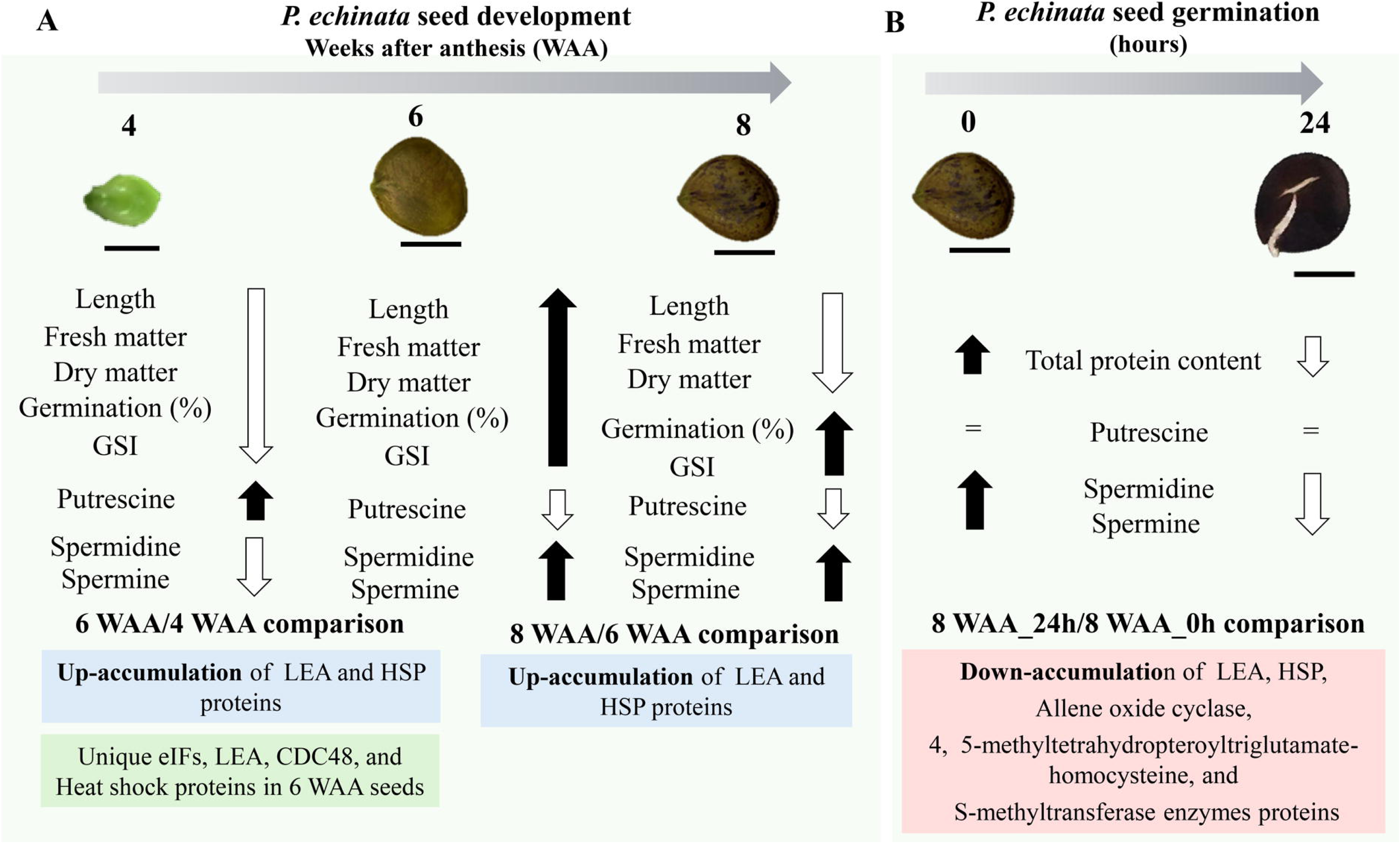
Schematic diagram showing the main developmental and biochemical changes in *P. echinata* seeds during development (A) and germination (B)

Unique and up-accumulated proteins identified in seeds of *P. echinata* at 6, compared to seeds at 4 WAA (Clusters 1 and 2), highlight an increased demand for energy and macromolecules. This demand supports essential processes required for seed growth, such as cell division, cell expansion, and the formation of specialized tissues for nutrient storage, as seeds progress toward maturation.^26^

At the beginning of protein synthesis, various eukaryotic translation initiation factors interact with methionyl tRNA, mRNA, and ribosomal subunits to facilitate the assembly of pre-initiation complexes.^63^ In *P. echinata* seeds, two eukaryotic translation initiation factor (eIF) proteins were up-accumulated at 6 compared to 4 WAA, and three were uniquely detected in seeds at 6 WAA. Among eIFs, eIF3 is the largest and most complex initiation factor, interacting with other eIFs and playing a crucial role in organizing proteins on the surface of the 40S ribosomal subunit.^63^ In addition to translation initiation, several ribosomal structural components were differentially regulated in seeds collected at 6 compared to those collected at 4 WAA, with 59 ribosomal proteins up-accumulated and 21 unique to seeds at 6 WAA. Ribosomal proteins play key roles in ribosome constitution, protein biosynthesis, transcription, translational regulation, and developmental processes in plants.^64^

Our results indicate that as protein synthesis increases, there is a corresponding increase in the demand for proteases that help control quality during seed development. Among them, the accumulation of ubiquitin-proteasome system (UPS)-related proteins in clusters 1 and 2 indicates an increase in proteolysis to maintain protein homeostasis during the development of *P. echinata* seeds. UPS is the main pathway for protein degradation in cells, regulating the abundance of key regulatory proteins and ensuring a healthy, properly functioning proteome.^65,66^

In *P. echinata*, 18 DAPs associated with proteasome activity were identified in the 6 WAA/4 WAA comparison, some of which were unique to seeds at 6 WAA, and some were up-accumulated in seeds at 6 compared to those at 4 WAA. In plants, the 26S proteasome regulates protein degradation through the ubiquitin/26S proteasome pathway, which plays a crucial role in development by influencing a wide range of processes, including embryogenesis and regulation of seed size and development.^65,67^ Moreover, seven homologs of CDC48 were differentially regulated in the 6 WAA/4 WAA comparison. In plants, CDC48 plays a regulatory role in development beyond protein degradation through the UPS and the endoplasmic reticulum-associated protein degradation (ERAD) system.^68^ In *A. thaliana*, CDC48 contributes to cell division and expansion, cytokinesis, and growth processes.^69^ Thus, the up- accumulation of UPS-related proteins and CDC48 proteins in seeds at 6 WAA compared to those at 4 WAA is related to the development of *P. echinata* seeds.

Another biological process enriched in Clusters 1 and 2 was a carbohydrate derivative metabolic process. Carbohydrate metabolism mainly includes glycolysis and the tricarboxylic acid (TCA) cycle, which provide most of the energy required for seed maturation and germination. In plants, the TCA cycle comprises enzymatic reactions that oxidize malate and pyruvate, producing NADH to generate ATP through the electron transport chain (ETC).^70^ Several enzymes associated with the TCA cycle were differentially regulated in the seeds of *P. echinata.* Among them, aconitate hydratase, which accumulates in seeds at 6 but not at 4 WAA, is an enzyme that catalyzes the reversible isomerization of citrate to isocitrate via cis-aconitate in the TCA cycle. In *A. thaliana*, the activity of the protein aconitate hydratase was shown to be essential for both embryo and seed development.^71^ Moreover, succinate-CoA ligase, up-accumulated in 6 compared to 4 WAA seeds and unique to 6 WAA seeds, catalyzes a reversible reaction that converts succinyl-CoA and ADP or GDP to succinate and ATP or GTP.^72^ The intermediates of the TCA cycle play crucial roles in providing carbon substrates for various biosynthetic pathways, including the synthesis of carbohydrates and fatty acids.^73,74^ On the other hand, malate synthase protein accumulates in 6 WAA seeds, but complements the reactions catalyzed by isocitrate lyase, which is unique to *P. echinata* seeds at 6 WAA, to produce malate and coenzyme A, allowing efficient utilization of two- carbon compounds derived from lipids, such as acetate, to generate energy.^75^

In *P. echinata*, key glycolytic proteins were up-accumulated or unique in seeds at 6 WAA compared to those at 4 WAA, suggesting that these proteins may also play a role in seed maturation and desiccation. You et al.^76^ correlated the increased accumulation of fructose-bisphosphate aldolase, phosphoglycerate kinase, glucose-6-phosphate isomerase, and glyceraldehyde-3-phosphate dehydrogenase in larger grains during rice grain filling. Similarly, the up-accumulation of these proteins in seeds at 6 WAA compared to 4 WAA may play a role in regulating size and influencing seed filling in *P. echinata*. In addition, phosphoglycerate kinases are involved in glycolysis/gluconeogenesis and photosynthetic carbon metabolism in plants.^77^ This protein converts 1,3-bisphosphoglycerate to 3-phosphoglycerate and ATP during glycolysis and participates in reverse-reaction gluconeogenesis and the Calvin-Benson cycle, providing intermediates for carbohydrate synthesis, such as starch and fatty acid biosynthesis.^77–79^ These findings suggest that energy production is crucial for the seed development process in *P. echinata*.

The up-accumulation of proteins associated with protein metabolic processes, including folding and refolding, in *P. echinata* highlights the importance of maintaining proper protein conformation and function during seed development. Many of these proteins were up-accumulated or unique to *P. echinata* seeds collected at 6 compared to those collected at 4 WAA (Cluster 2), with four disulfide isomerase proteins and 22 HSPs. HSPs expressed during seed maturation not only play a critical role in response to environmental stresses but also serve as chaperones to facilitate the assembly of newly synthesized proteins during developmental transitions.^80^ Initially, detected in response to abiotic stresses, sHSPs are also produced in reproductive organs during developmental processes such as seed maturation, germination, and fruit maturation, even in the absence of stress.^81,82^

In *P. echinata,* 15 LEA proteins were differentially accumulated. Although these proteins were not specifically enriched in any particular biological process, the greater accumulation or unique of 14 LEA proteins in seeds collected at 6 than in those collected at 4 WAA (Clusters 1 and 2), as well as the up- accumulation of six LEA proteins in both 6 WAA/4 WAA and 8 WAA/6 WAA comparisons, suggest the relevance of this type of protein in the maturation and desiccation phases in seeds of *P. echinata.* At this stage, as orthodox seeds acquire the ability to withstand dehydration, LEA proteins are associated with desiccation tolerance acquisition.^83^

The major biological process enriched in Cluster 3 was photosynthesis. The reduction in the accumulation of proteins involved in photosystems in seeds at 6 compared to those of 4 WAA (Cluster 3) indicates that the seeds of *P. echinata* are dependent on cytosolic processes to provide ATP, reducing power, and carbon precursors, which are required for seed development and the biosynthesis of storage reserves.

Another biological process enriched in Cluster 3 of *P. echinata* seeds was the flavonoid biosynthetic process, with a reduction in the accumulation of several proteins related to this pathway in 6 WAA seeds compared to those of 4 WAA seeds. Flavonoids, as a diverse class of phenolic compounds, play critical roles in plant environment interactions, pigmentation, defense against biotic and abiotic stresses, and various physiological or developmental functions, including seed physiology.^84^ However, as seeds progress to more advanced stages of development (6 WAA), the reduction in the accumulation of these proteins suggests a metabolic shift prioritizing the deposition of energy reserves, such as lipids and proteins, which are essential for the development of *P. echinata* seeds.

Our findings also highlight a correlation between specific proteins and PAs in *P. echinata* seeds. Among these proteins, seven CDC48 homologs were differentially accumulated, with six being unique to seeds at 6 WAA and one being up-accumulated in 6 WAA seeds compared to 4 WAA seeds. CDC48 is known for its role in cell division, cytokinesis, and cell expansion. However, a potential regulatory function of CDC48 in the synthesis and metabolism of S-adenosylmethionine (SAM) has been proposed.^68^ Four proteins associated with methionine synthesis, a precursor of SAM, had greater accumulation in 6 WAA seeds than in 4 WAA seeds (Cluster 2). Among them are two adenosylhomocysteinases and two 5-methyltetrahydropteroyltriglutamate-homocysteine S-methyltransferases. In plants, 5-methyltetrahydropteroyltriglutamate-homocysteine methyltransferases play crucial roles in the methionine biosynthesis pathway, catalyzing the methylation of homocysteine to methionine through the use of 5-methyltetrahydrofolate as a methyl donor.^85^ Conversely, adenosylhomocysteinases are responsible for breaking down S-adenosylhomocysteine (SAH), a byproduct of SAM-dependent methylation reactions, into adenosine and homocysteine.^86^ Methionine serves not only as a crucial amino acid for protein synthesis but also as a precursor for SAM, a universal methyl donor involved in various metabolic pathways, including PA biosynthesis.^87^ In *P. echinata* seeds, five eIFs were DAP in the comparison of 6 WAA and 4 WAA seeds. eIF5A activity is crucial during the early stages of embryo development and processes triggered after fertilization, underscoring its importance in plant development.^88^

Our findings indicate a correlation between Put and the immature state of the seeds, as well as the absence of germination in *P. echinata* seeds harvested at 4 WAA, a stage in which cell division is more intense.^17,89^ In contrast, the higher contents of Spd and Spm observed during the later stages of *P. echinata* seeds (6 and 8 WAA) highlight the relevant role of Spd and Spm during the maturation and desiccation phases. The relatively high contents of Spd and Spm observed in the latter stages of development in several species are associated with seed maturation, when growth results predominantly from cell elongation and accumulation of storage reserves.^17,90^

### 4.3. Reduction in the abundance of some proteins and PAs is crucial for the successful germination of *P. echinata* seeds

Seed germination is a vital stage in plants, during which cells from embryos undergo a programmed transition from a quiescent state to an active metabolic state.^91^ The progressive increase in the percentage of germination and the GSI with seed maturation indicates that seeds at 4 WAA are immature and require an additional two weeks to reach the germination potential. Consequently, the increase in both the percentage of germination and GSI reflects the physiological maturation of the seeds, with maximum values observed in those harvested at 8 WAA.

During the seed germination process of *P. echinata*, a greater number of down-accumulated than up-accumulated proteins, including LEA and HSP proteins, was detected in 8 WAA seeds at 24 h after imbibition than in 8 WAA seeds before imbibition (8 WAA_24h/8 WAA_0h comparison). Upon imbibition, the stored molecules participate in the germination process at various levels. LEA proteins, which play a critical role during seed maturation and desiccation, are known to degrade during the first 24 h of imbibition.^92^ In *P. echinata*, in addition to the increased abundance of several LEA proteins during seed development (Clusters 1 and 2), four LEA proteins showed decreased abundance within the first 24 h of germination. This observed decrease in the abundance of LEA proteins may facilitate the resumption of metabolic activities essential for germination in this species. Furthermore, two of the identified enriched biological processes were related to protein folding and response to stress, and most proteins decreased abundance during imbibition, including two HSPs. Sung et al.^93^ reported high levels of preserved Hsp70 transcripts in mature dormant *Arabidopsis* seeds, which decreased rapidly within the first 12 h of imbibition. Similarly, HSP proteins were detected only during the early stages of seed germination in *C. legalis*.^29^ Thus, the reduction in the accumulation of proteins related to protein folding and the response to stress throughout seed germination of *P. echinata* may be a more relevant process for rehydration and energy mobilization than the accumulation of new proteins.

Additionally, proteins related to enriched processes, small molecule metabolic process, carboxylic acid metabolic process, and organic substance biosynthetic process decrease in abundance during *P. echinata* seeds during germination. This reduction in accumulation suggests a metabolic change after imbibition, where priority transitions from reserve storage, stress tolerance, and biosynthetic activities to active growth and cellular rehydration in seeds of *P. echinata* 24 h after imbibition. Studies support this dynamic regulation, highlighting that these processes are tightly controlled during germination to ensure the mobilization of stored metabolites for energy and biosynthetic needs.^92^ Among the proteins that showed decreased abundance in these biological processes is allene oxide cyclase 4, which is involved in jasmonic acid (JA) biosynthesis,^94^ that has been shown to inhibit seed germination in several species.^95^

Furthermore, two 5-methyltetrahydropteroyltriglutamate-homocysteine S-methyltransferase enzymes, associated with carboxylic acid metabolism, biosynthesis of small organic molecules, and broader organic biosynthetic processes, were reduced in abundance in 8 WAA seeds 24 h after imbibition compared to before. As mentioned previously, this enzyme plays a crucial role in the methionine biosynthesis pathway, which serves as a precursor for SAM, which is necessary for Spd and Spm synthesis.^85^ In this sense, the reduction in the contents of Spd and Spm during germination can be related to the reduction in the accumulation of this protein in 8 WAA seeds of *P. echinata* 24 h after imbibition.

The homeostasis of PA in plants is regulated through the modulation of biosynthesis, transport, conjugation, interconversion, and catabolism.^87,96^ Maintaining the PA content is necessary to avoid depletion or overaccumulation, which can alter cell viability in various organisms.^87^ In *P. echinata*, the fact that the Put content remained constant during seed imbibition and germination suggests that there was no degradation or interconversion of this PA into Spm and Spd.

Our findings reveal that seed germination in *P. echinata* involves tightly coordinated molecular processes to support growth, energy production, and stress management through a decrease in the abundance of some proteins, as well as PAs, that accumulate during seed development. Therefore, the patterns of protein and PA changes during germination demonstrate the intricate balance between biosynthetic activity, stress management, and reserve mobilization, which are crucial for the successful germination of *P. echinata* seeds (Figure 11B).

## 5. Conclusions

This study provides novel insights into the molecular mechanisms underlying seed maturation and germination in *P. echinata*. The development of a nonredundant, species-specific protein databank significantly improved protein identification compared to the use of the *G. max* database, and the sequential protein extraction method proved highly effective for proteomic analysis. During early seed development, translation-related proteins and proteasome components were highly accumulated, whereas stress-associated proteins, including LEA and HSPs, peaked during seed maturation. The abundance of carbohydrate metabolism-related proteins highlights their role in providing energy reserves for germination. Proteins involved in methionine biosynthesis appear to mediate the regulatory functions of polyamines, with putrescine associated with early development and spermidine and spermine associated with maturation, tolerance to desiccation, and germination. In addition, reduction in certain proteins and polyamines that accumulated during seed development appears necessary for successful germination. In general, these findings advance our understanding of seed biology in *P. echinata* and can inform future research on conservation and propagation strategies, including somatic embryogenesis, for this endangered species.

## Authors’ contributions

Claudete Santa-Catarina conceived and supervised the study, acquired funding, and contributed to methodology, project administration, and manuscript review and editing. Rosana Gobbi Vettorazzi contributed to the conceptualization, methodology, formal analysis, investigation, data curation, visualization, and wrote the original draft. Vanildo Silveira contributed to proteomic methodology, proteomic data collection, and manuscript review and editing. Kariane Rodrigues de Souza conducted proteomic investigations. Tadeu dos Reis de Oliveira contributed to proteomic data collection and visualization.

Clicia Grativol, Geovanna Vitória Olimpio, and Vitor Batista Pinto contributed to the RNA extraction methodology. Thiago Motta Venancio and Gabriel Quintanilha-Peixoto performed RNA-seq data collection, software and bioinformatics analyses, and contributed to database generation. All authors read and approved the final manuscript.

## Acknowledgments

This work was supported by the Conselho Nacional de Desenvolvimento Científico e Tecnológico (CNPq) [grant numbers 309303/2019-2, 312595/2023-9]; the Fundação Carlos Chagas Filho de Amparo à Pesquisa do Estado do Rio de Janeiro (FAPERJ) [grant numbers E26/202.533/2019, E26/210.088/2022, E26/200.396/2023]; and the Coordenação de Aperfeiçoamento de Pessoal de Nível Superior – Brasil (CAPES) [Finance Code 001]. The sponsors had no role in study design, data collection and analysis, decision to publish or manuscript preparation. We also thank the Life Sciences Core Facility (LaCTAD) staff at the State University of Campinas (UNICAMP), Campinas, SP, Brazil, for their assistance with RNA sequencing analysis.

## Declaration of Competence of Interest

The authors declare that they have no known competing financial interests or personal relationships that could have appeared to influence the work reported in this paper.

## Data availability

The mass spectrometry proteomics data and the nonredundant protein database of *P. echinata,* generated from RNA-seq data, have been deposited in the ProteomeXchange Consortium through the PRIDE partner repository with the dataset identifier PXD056397. All data are available from the corresponding author

## Abbreviations

DAP –: differentially accumulated protein
DM -: Dry matter
eIF -: Translation initiation factor
FM -: Fresh matter
GSI -: Germination speed index
HSP -: Heat shock protein
LEA -: Late embryogenesis abundant
PA –: Polyamine
PPI -: Protein-protein interaction
Put -: Putrescine
SAM -: S-adenosylmethionine (SAM)
Spd -: Spermidine
Spm –: Spermine
TCA -: Tricarboxylic acid cycle
UPS -: Ubiquitin-proteasome system
WAA -: Weeks after anthesis

## Supplementary tables

**Table S1.** Complete list of all proteins identified in *Paubrasilia echinata* during seed development, using the *P. echinata* generated databank. Proteins with significant results in the Student’s t-test (two-tailed; P < 0.05) were classified as DAPs, with up-accumulated proteins defined by a log2 fold change (FC) greater than 0.6, and down-accumulated proteins defined by a log2 FC less than −0.6.

**Table S2.** Complete list of all proteins identified in *Paubrasilia echinata* during seed development, using the UNIPROT *Glycine max* databank. Proteins with significant results in the Student’s t-test (two-tailed; P < 0.05) were classified as DAPs, with up-accumulated proteins defined by a log2 fold change (FC) greater than 0.6, and down-accumulated proteins defined by a log2 FC less than −0.6.

**Table S3**. Complete list of all proteins identified in *Paubrasilia echinata* during seed germination, using the *P. echinata* generated databank. Proteins with significant results in the Student’s t-test (two-tailed; P < 0.05) were classified as DAPs, with up-accumulated proteins defined by a log2 fold change (FC) greater than 0.6, and down-accumulated proteins defined by a log2 FC less than −0.6.

**Table S4.** Complete list of all proteins identified in *Paubrasilia echinata* during seed germination, using the UNIPROT *Glycine max* databank. Proteins with significant results in the Student’s t-test (two-tailed; P < 0.05) were classified as DAPs, with up-accumulated proteins defined by a log2 fold change (FC) greater than 0.6, and down-accumulated proteins defined by a log2 FC less than −0.6.

**Table S5.** BLAST results from the alignment between proteins identified using both Glycine max and *Paubrasilia echinata* databanks during seed development.

**Table S6.** BLAST results from the alignment between proteins identified using both Glycine max and *Paubrasilia echinata* databanks during seed germination.

**Table S7.** STRING enrichment results for biological processes at the protein cluster level in identified proteins of *Paubrasilia echinata* during seed development.

**Table S8.** STRING enrichment results for biological processes in identified proteins of *Paubrasilia echinata* during seed germination.

---

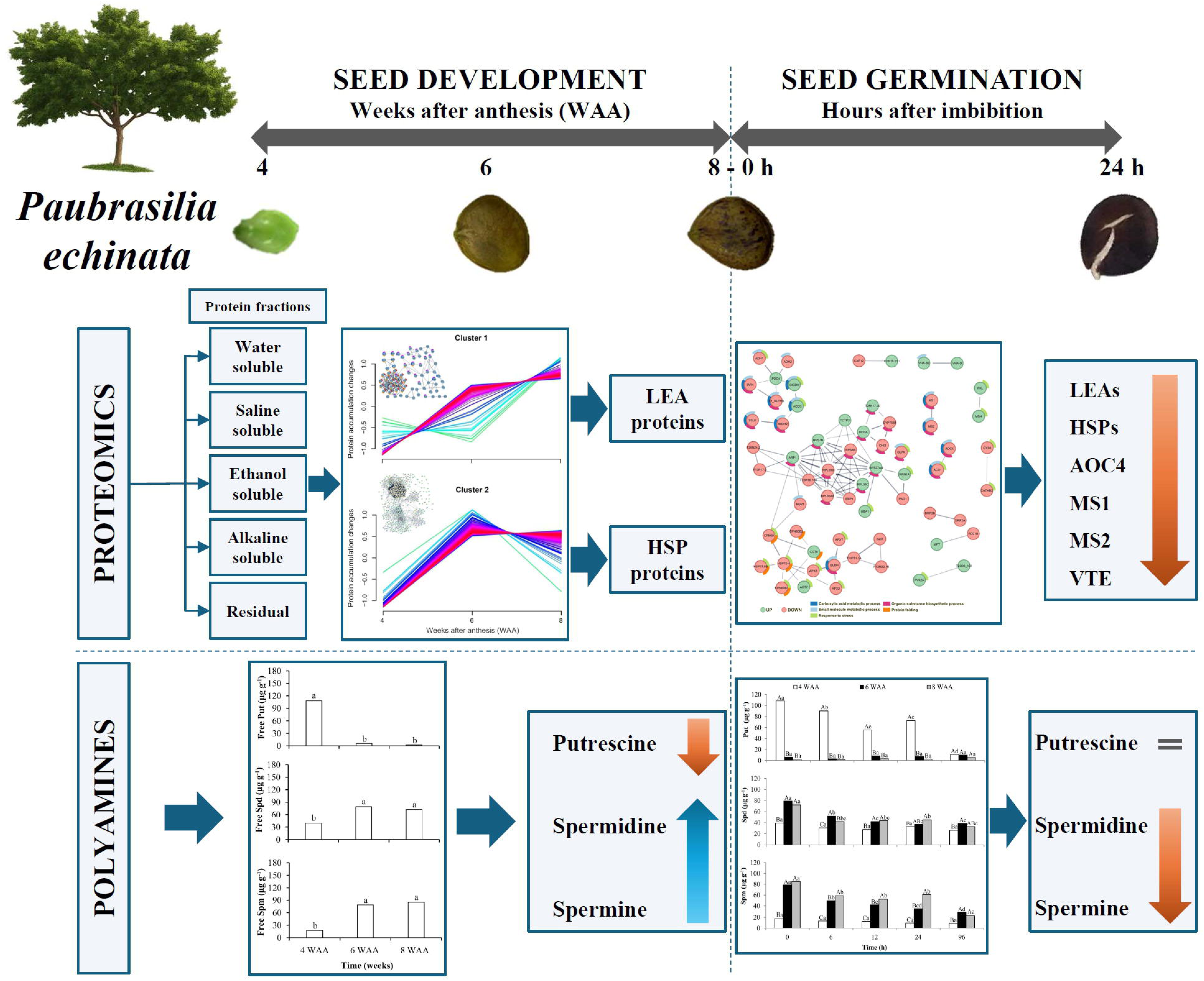

